# Oncogenic G12D mutation alters local conformations and dynamics of K-Ras

**DOI:** 10.1101/178483

**Authors:** Sezen Vatansever, Burak Erman, Zeynep H. Gümüş

**Author notes:** Corresponding authors: (BE) and (ZHG).

## Abstract

K-Ras is the most frequently mutated oncoprotein in human cancers, and G12D is its most prevalent mutation. To understand how G12D mutation impacts K-Ras function, we need to understand how it alters the regulation of its dynamics. Here, we present local changes in K-Ras structure, conformation and dynamics upon G12D mutation, from long-timescale Molecular Dynamics simulations of active (GTP-bound) and inactive (GDP-bound) forms of wild-type and mutant K-Ras, with an integrated investigation of atomistic-level changes, local conformational shifts and correlated residue motions. Our results reveal that the local changes in K-Ras are specific to bound nucleotide (GTP or GDP), and we provide a structural basis for this. Specifically, we show that G12D mutation causes a shift in the population of local conformational states of K-Ras, especially in Switch-II (SII) and α3-helix regions, in favor of a conformation that is associated with a catalytically impaired state through structural changes; it also causes SII motions to anti-correlate with other regions. This detailed picture of G12D mutation effects on the local dynamic characteristics of both active and inactive protein helps enhance our understanding of local K-Ras dynamics, and can inform studies on the development of direct inhibitors towards the treatment of K-Ras^G12D^-driven cancers.

## Introduction

K-Ras is the most frequently mutated oncoprotein in multiple human cancers ^1-3^. Patients with oncogenic K-Ras mutations have very poor response to standard therapies. Unfortunately, these mutations eventually emerge during the course of their treatment and drive resistance ^4-6^. The importance of oncogenic mutations in K-Ras stem from its pivotal role in signaling networks that control cellular growth, proliferation, and differentiation ^7^. To perform its cellular roles, K-Ras continuously switches between GDP-bound (inactive) and GTP-bound (active) states (Fig 1A) ^8,9^. This switch mechanism is important in turning signals through K-Ras on or off, because only active K-Ras can bind to and trigger its downstream proteins ^10^. During the switch, active K-Ras (K-Ras-GTP) acts as a GTPase by hydrolyzing GTP and becomes inactive (K-Ras-GDP). K-Ras-GTP may also bind to GTPase-activating proteins (GAPs) that accelerate this process ^11,12^. However, oncogenic K-Ras mutations impair both its GTPase activity and GAP protein binding, inhibiting GTP hydrolysis. Thus, unable to switch to its GDP-bound (inactive) state, mutant K-Ras remains continuously active, leading to prolonged activation of its downstream pathways associated with oncogenic cellular growth ^10, 13-15^. There is strong evidence that blocking mutant K-Ras activity can be very effective in treatment ^16,17^. Yet, despite decades of research, there are still no drugs in the clinic today that directly target K-Ras mutants ^18,19^.

**Figure 1.**
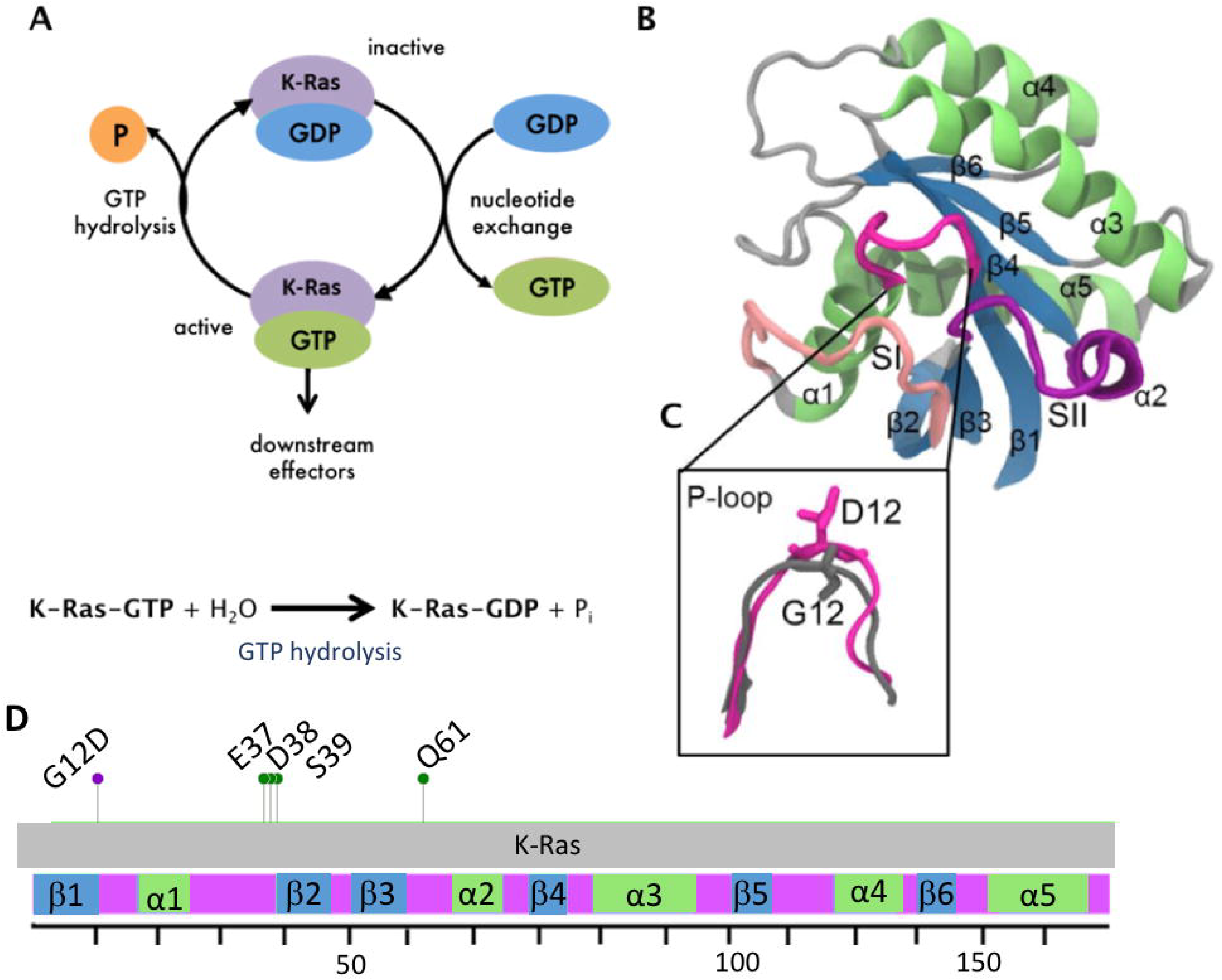
K-Ras switch mechanism and 3D structure. (A) Cartoon representation of K-Ras switch mechanism between GTP-bound (active) to GDP-bound (inactive) forms and the GTP hydrolysis reaction (B) Cartoon representation of K-Ras protein consisting of secondary structures. Blue: β-sheets, green: α-helices, pink: P-loop, light pink: SI region, purple: SII region. (C) Zoomed-in visualization of the P-loop. Grey: Wild-type, Pink: G12D mutant. Residues G12 and D12 are in stick representation. (D) G12D mutation site (red), protein-protein interaction site residues for its regulator and downstream effector proteins (green) and schematic representation of K-Ras sequence (residues 1-169).

Activating oncogenic K-Ras mutations are frequently observed at residue positions 12, 13 and 61 in cancer patients. Among these, G12 is the most frequently mutated residue (89%), and it most often mutates to aspartate (G12D, 36%) followed by valine (G12V, 23%) and cysteine (G12C, 14%) ^3,10^. G12 is located at the protein active site, which consists of a phosphate binding loop (P-loop, residues 10-17) and two switch regions (SI, residues 25-40, and SII, residues 60-74) (Fig 1B-C). The residues in the active site bind to the phosphate groups of GTP and are responsible for the GTPase function of K-Ras. The switch regions SI and SII are additionally responsible for controlling binding to effector and regulator proteins (Fig 1D). However, the mutation of glycine at position 12 to aspartate (G12D) leads to the projection of a bulkier and negatively charged side group into the active site, which causes a steric hindrance in GTP hydrolysis ^20^, impairs the GTPase function and locks K-Ras in its active (GTP-bound) state ^12^. NMR studies have shown that Ras proteins acquire a range of conformations during their motions in this GTP-bound form ^21-23^. Identifying the changes in dynamic conformations of K-Ras upon G12D mutation requires an integrated analysis that targets local conformational and dynamic changes at an atomistic level. Although the effects of G12D mutation on the structure, conformation and flexibility of K-Ras have been studied ^24-27^, how it alters the balance of local conformational states, and thereby the local dynamics of K-Ras still remains to be understood. This is an important question, as there is increasing evidence that crystal structure studies alone may miss drug-binding pockets on mutant K-Ras surface, while studies that include dynamics information have recently achieved promising results ^28-31^. However, these studies are so far limited to G12C mutant and local dynamic changes remain unknown. While it is plausible that the development of targeted drugs may remain elusive, understanding the unique dynamic characteristics of the most common oncogenic mutant K-Ras, G12D, can inform research efforts towards this goal.

Here, we present a computational procedure that reveals how the most prevalent oncogenic K-Ras mutation, G12D, triggers structural, conformational and dynamic changes that result in constitutive activation of the protein. Our integrated analysis particularly targets the local changes in the protein structure, conformation and dynamics, since global metrics are mainly affected by loop motions (i.e. rotations and translations) that are not related with its function ^32^. We are motivated by recent studies that have utilized protein dynamics data successfully to understand the effects of mutations ^33-35^. Most notably in drug discovery, dynamics data on oncogenic proteins ^25, 36-39^ have helped identify cryptic or allosteric binding sites ^40-43^. Encouraged by these results, we have hypothesized that the mutation-specific dynamic behavior of active and inactive K-Ras can best be explored by detailed analyses of their MD simulation data, from which we can investigate the range of their dynamic conformations and their atomistic-scale structural basis. To test our hypothesis, we used an integrated computational analysis that quantifies mutation-based local changes in protein conformations and their dynamic consequences.

In summary, we have performed replicate long time MD simulations (1 microsecond) of both wild-type and G12D mutant K-Ras (K-Ras^G12D^) in GTP-bound (active) and GDP-bound (inactive) forms. Briefly, we first studied the changes in the population of local conformational states caused by the mutation by evaluating the alterations in residue pair distances and local volume. Then, we explored the residue-specific population of conformational states of K-Ras and the population shift upon G12D mutation, and elucidated the structural changes that alter the formation of H-bonds and salt bridges that govern population balance of local conformational states. We then identified changes in local dynamics through a multi-step process where we quantified the fluctuations of all residues; correlated fluctuations for all residue pairs; and identified lost or newly formed correlations upon mutation. Finally, we related the observed structural changes to conformational and dynamic alterations, which enabled us to decode the important local changes that affect K-Ras function due to the G12D mutation. Overall, our study enhances our understanding of K-Ras^G12D^ dynamics and can inform studies on the development of direct inhibitors.

## Results

### Residue pair distance calculations show that G12D mutation causes local conformational changes in K-Ras

To understand the local conformational changes around each residue upon G12D mutation, we first analyzed the molecular dynamics (MD) simulation data of both active (GTP-bound) and inactive (GDP-bound) forms of wild type and G12D mutant K-Ras by using conventions from the Gaussian Network Modeling (GNM) approach, a widely employed tool in the analysis of protein dynamics. Specifically, we defined *the first coordination shell* of a residue as a sphere of radius ∼7.2Å ^44-47^, and the *second coordination shell* as twice the volume of the first, at a radius of ∼9.1Å ^47^ (See Methods). We then compared the distances between all residue pairs within *their second coordination shells* in wild-type and vs. mutant K-Ras. For this purpose, we represented the changes in the time-averaged distance between two residues *i* and *j* that are within their second coordination shells by Δ*R̄_ij_* = *R̄_ij,mutated_* = *R̄_ij,WT_* where *R̄_ij_* is the time averaged distance between residues *i* and *j*. In Fig 2A, we show the ∆*R̄_ij_* values for all residue pairs, where K-Ras ^WT^ is the reference and K-Ras^G12D^ is the final structure. As seen from the abundance of positive ∆*R̄_ij_* values from Fig 2A, the dominant distortion of the protein upon mutation is expansion. Specifically, i) the SII loop (Q61-S65) moves away from α3 (D92-Q99); ii) residues A11-G13 (P-loop) move away from Q61H (SII) and iii) the C-terminal residue of the α2 helix (M72) move away from the N-terminal residues of the same helix (S65, R68) and α3 (Q99). On the other hand, we observed that some residue pairs got (weakly) closer in K-Ras^G12D^, including D69 (SII)-V103 (α3) and L23 (α1)-V152, F156 (α5), demonstrated by their negative ∆*R̄_ij_* values in Fig 2A.

**Figure 2.**
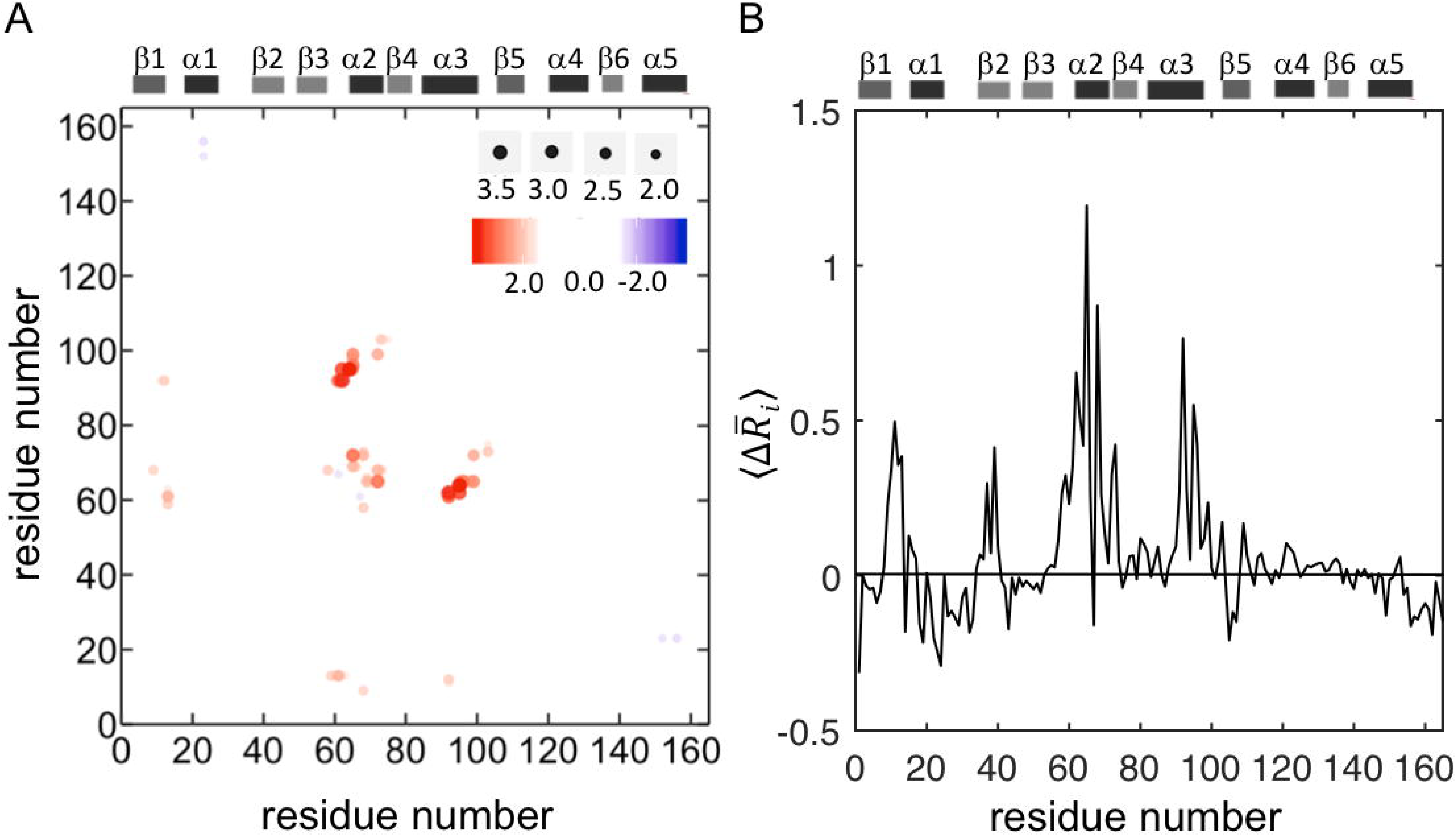
Conformational changes in K-Ras upon G12D mutation. (A) The differences in average pairwise distances (Δ*R̄_ij_*) where K-Ras^WT^ is the reference state. Red dots (positive Δ*R̄_ij_* values) show that pairs move further apart and blue dots (negative Δ*R̄ij* values) show that pairs move closer in K-Ras^G12D^. (B) The average of all Δ*R̄_ij_* values for each residue, 〈 Δ*R̄i*〉. The initial state is K-Ras^WT^ and the final state is K-Ras^G12D^. The residues which move away from their neighbors have positive values and dominate the mutant protein; residues that move close to their neighbors have negative values.

### G12D mutation alters the distribution of conformations of active K-Ras

To better understand the conformational changes upon G12D mutation, we explored the distribution of distances between all residue pairs for both active (GTP-bound) and inactive (GDP-bound) states of K-Ras (Fig 2A) from their MD simulation data. We observed that the wild type and mutant proteins exhibited distinct distribution patterns. Specifically, while the residue pair distances in both inactive (GDP-bound) wild-type and mutant K-Ras exhibited broad distributions with multiple peaks (Fig 3-4), in active (GTP-bound) K-Ras, they exhibited narrow distributions in wild-type with two distinct peaks and only one peak in mutant.

**Figure 3.**
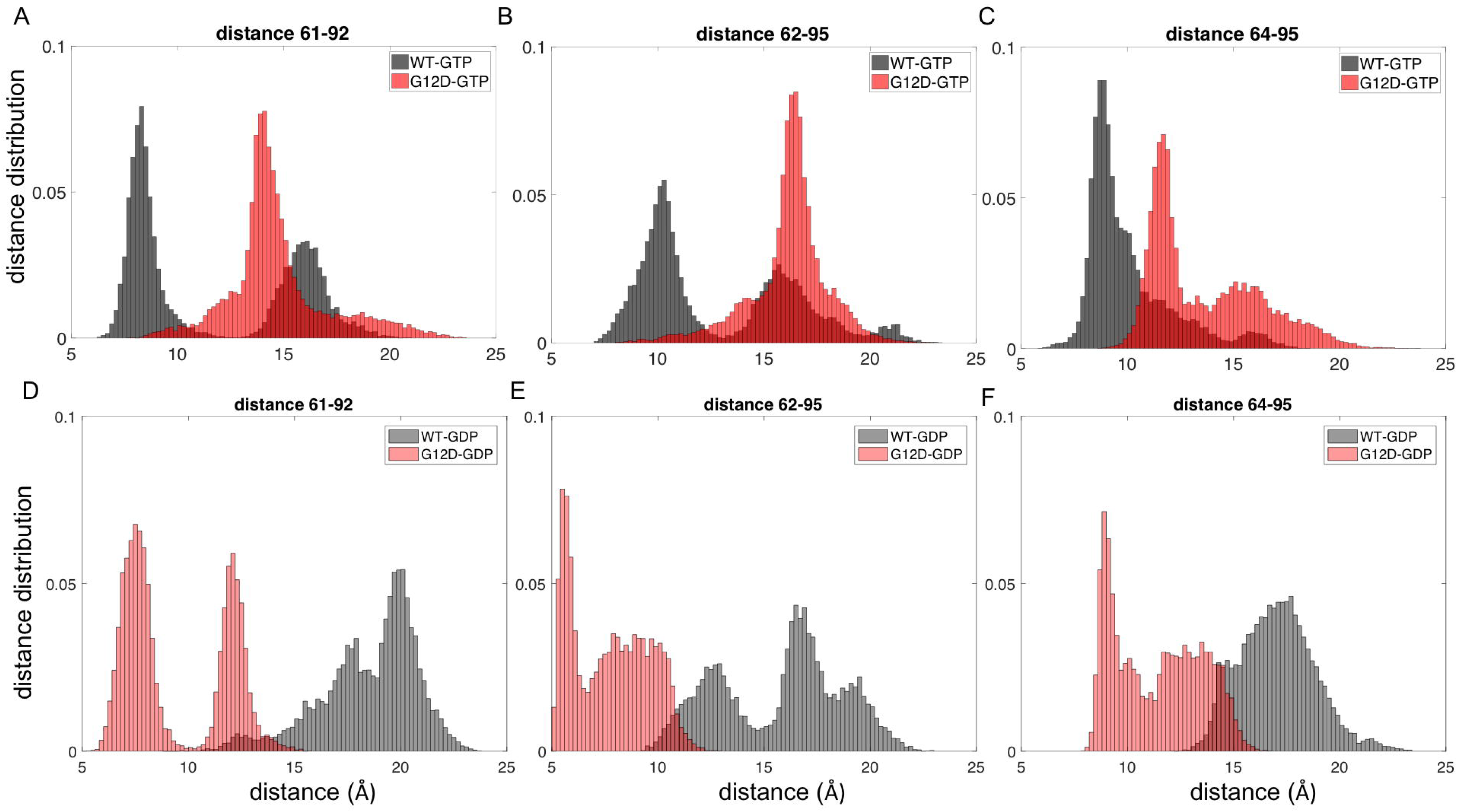
Distribution of distances between residue pairs in SII-α3 region. Distance distributions of Cα of residue pairs in K-Ras^WT^-GTP (black) and K-Ras^G12D^-GTP (red); (A) Q61-D92, (B) E62-H95; (C) Y64-H95; in K-Ras^WT^-GDP (grey) and K-Ras^G12D^-GDP (pink); (D) Q61-D92; (E) E62-H95, (F) Y64-H95.

**Figure 4.**
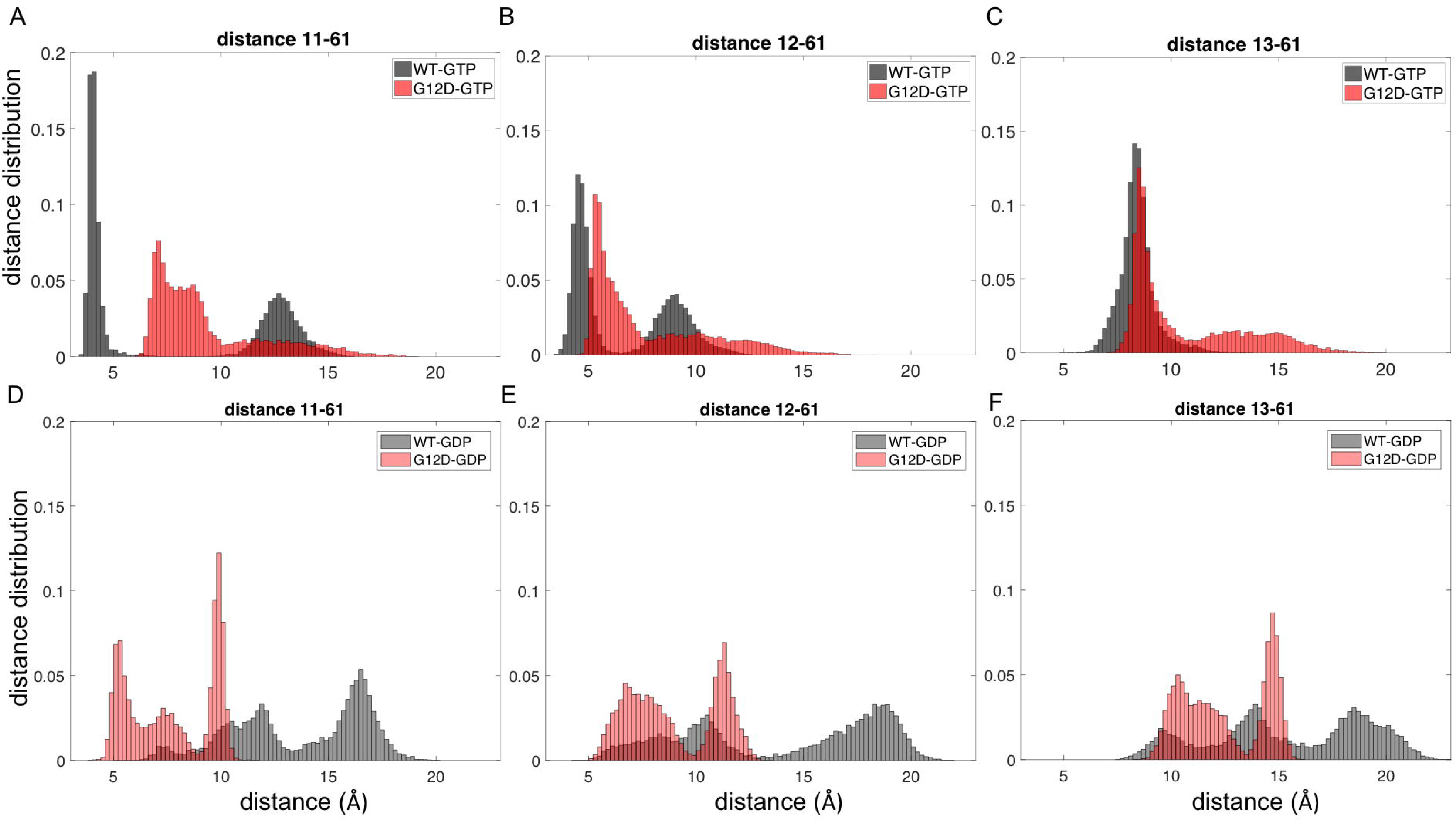
Distribution of distances between residue pairs in the P loop-SII region. Distance distributions between Cαs of residue pairs in K-Ras^WT^-GTP (black) and K-Ras^G12D^-GTP (red) (A) A11-Q61, (B) G12D-Q61, (C) G12D-Q61; in K-Ras^WT^-GDP (grey) and K-Ras^G12D^-GDP (pink) (D) A11-Q61, (E) G12D-Q61, (F) G12D-Q61.

### G12D mutation alters the balance of local conformational states between SII and α3 regions

Based on our analysis of all residue pairs, the largest conformational change due to G12D mutation in active K-Ras was between SII and α3 regions. We observed that the distances between residue pairs 61-92, 62-92, 62-95 and 64-95 populate multiple conformations in inactive (GDP-bound) K-Ras^WT^ (Fig 3). However, GTP binding causes these pairs to have two distinct conformations, where the peak of the distance distribution of the first conformation is at a smaller distance and the peak of the second conformation is at a larger distance (Fig 3A-C). However, G12D mutation decreases the number of conformations of the active protein, and the residue pairs assume only one conformation, which is similar to the second conformation of active wild type. Interestingly, the mutation affects the GDP-bound protein differently: while the distances between the residues still populate multiple conformations as in GDP-bound K-Ras^WT^, peaks are at shorter distances, and are in fact similar to the first conformation of GTP-bound K-Ras^WT^ (Fig 3D-F).

### G12D mutation driven changes in conformational states between SII and α 3 regions are due to the brakeage of H-bonds and the formation of a salt bridge between them

SII region consists of the SII loop (A59-Y64, N-terminus) and the α2 helix (residues S65-T74, C-terminus). In wild-type K-Ras-GTP, α2 helix of SII interacts with α3 helix through an H-bond between R68 and D92 in mutant K-Ras-GTP and in the second conformational state. Because of the omega shape of the SII loop, the N-terminal residues of SII (Q61, E62 and Y64) assume a distant conformation from α3 helix at the presence of the H-bond between α2-α3, while the C-terminal residues of SII move toward α3 (D69-V103 pair in Fig 5). In the first conformational state of wild type K-Ras-GTP, this H-bond is not present and thus E62 on the SII loop forms a salt bridge with K88, while K88 forms a salt bridge with D92. The SII loop and α3 helix get closer via these salt bridges. Similarly, in mutant K-Ras-GDP, the formation of an H-bond between the SII loop and α3 helix regions drives these regions to switch between two conformations: a close conformation through the H-bond of E63 (SII)-Q99 (α3) and an open conformation in its absence. However, in wild-type K-Ras-GDP simulations, we did not observe the E63-Q99 H-bond, and the SII loop-α3 residue pairs were further apart.

**Figure 5.**
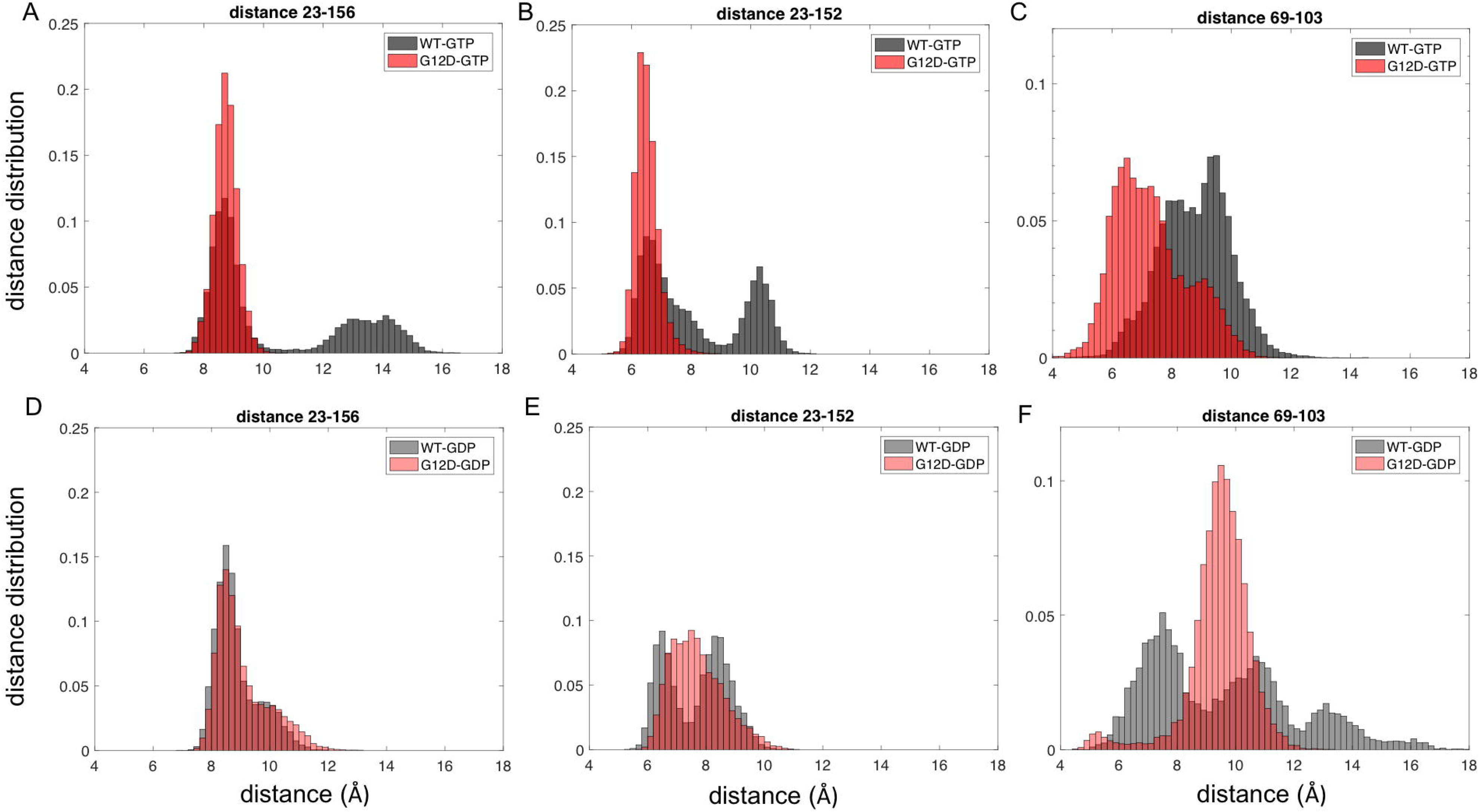
Distance distributions of residue pairs which get closer after G12D mutation in K-Ras-GTP in K-Ras^WT^ (black) and K-Ras^G12D^ (red). Distance distribution of Cα of residue pairs in K-Ras^WT^-GTP (black) and K-Ras^G12D^-GTP (red) (A) L23-V152, (B) L23-F156, (C) D69-V103; in K-Ras^WT^-GDP (grey) and K-Ras^G12D^-GDP (pink) (D) L23-V152, (E) L23-F156, (F) D69-V103.

### G12D mutation causes a population shift in the conformations of Q61-P-loop

For inactive (GDP-bound) K-Ras^WT^, the distance distribution plots between P-loop residues A11-G13 and SII residue Q61 (Fig 4) show multimodal distributions, indicating three different conformational states (Fig 4D-F). However, in active (GTP-bound) K-Ras^WT^ plots (Fig 4A-B), A11-Q61 and G12-Q61 pairs show only two distinct conformational states, which are similar to those between SII-α3 regions. The first conformation peak is close to an H-bond distance, while the second conformation peak shifts to a longer distance. In contrast, the distances between G13-Q61 show normal distribution with one peak at a longer distance (Fig 4C). This different conformation of G13 relative to Q61 may have resulted from the omega shape of the P-loop, which turns at the C-terminal neighborhood of G12. After G12D mutation, the number of conformations of the pairs A11-Q61 and D12-Q61 decrease. In active (GTP-bound) mutant K-Ras, these two pairs, in addition to G13-Q61, exhibit broad distance distributions with one peak (Fig 4A-C). Upon inactivation of mutant K-Ras, these three pairs can obtain multiple conformations similar to the inactive wild type K-Ras as seen in Figure 4D-F.

### G12D mutation leads to the formation of a permanent salt bridge between the P-loop and SII regions in active protein

In wild type K-Ras-GTP simulations, we observed that a salt bridge between K16 (α1) and E63 (SII) leads to two distinct conformations of the P-loop-SII pairs. Residue K16 resides at the C-terminal end of the P-loop, while the residues A11 and G12 reside at its N-terminus. When K16 forms the salt bridge with E63 (SII), A11-G12 get distant from Q61 (SII) corresponding to the second confirmation in Figure 4, and when this salt bridge dissappers, A11-G12 get closer to Q61 as in the first conformation. However, in mutant K-Ras-GTP simulations, this salt bridge does not disappear. Furthermore, K16 forms an H-bond with G10 on the P-loop (note that the mutated residue at position 12 is also on this P-loop). Because of this H-bond, A11-D12 and Q61 pairs are located at a distance between the first and the second conformations of A11-G12 and Q61 pairs in wild-type (Fig 4A-C).

### G12D mutation causes α5 helix to move towards α1 helix in active K-Ras

In addition to the residue pairs that assume distant conformations after mutation, we also investigated the distance distributions of the α1 and α5 regions, and observed that they got closer in active K-Ras^G12D^ (Fig. 5). Specifically, we observed that in active K-Ras^WT^, the residue pair distance distribution curves for L23 (α1)-V152 (α5) and L23 (α1)-F156 (α5) switched between a closer (6.5 Å for L23-V152, 8.5 Å for L23-F156) and a distant (10.5 Å for L23-V152, 14 Å for L23-F156) conformational state. Furthermore, the inactive wild-type and inactive mutant proteins, had similar patterns in their distribution curves for these residue pairs. However, in the active K-Ras^G12D,^ the distance distribution curves of these pairs had single peaks (6.5 Å for L23-V152, 8.5 Å for L23-F156), similar to their closer conformational states in active K-Ras^WT^.

### G12D mutation leads to changes in conformations of residues relative to their neighborhood

According to GNM, a residue typically fluctuates within its first or second coordination shells ^44,45^. Within these volumes, there are several other residues, which are either near-neighbors along the chain or are spatially distant. As has been shown by the GNM model, ^45^, a residue with a smaller number of neighbors will show larger fluctuations than a residue with a larger number of neighbors. Therefore, the neighborhood of a given residue significantly affects its fluctuations and dynamics. To understand which parts of K-Ras move away from its neighbors and which parts move closer upon mutation globally, we calculated the average of all ∆*R̄_ij_* values (the time averaged distance between two residues i and j) for each residue *i*, 〈 ∆*R̄_ij_*〉. Overall, we observed that after the G12D mutation, most protein parts, especially the P-loop, SI, SII and α3, move away from their neigbors, suggesting larger fluctuations (Fig 2B).

We, then, aimed to understand the relation between the changes in conformations of a residue pair and the changes in individual conformations of each residue in that pair. In the distance distribution calculations (Fig 3-5), we present the results of the residue pairs that underwent the largest change in distance due to G12D mutation. For each residue in the identified pairs, we estimated the extent of its deviation from its neighbors by comparing the individual ∑_*j*_ ∆*R̄_ij_*, values of the residues in the identified pairs (Supplementary Table S1). We discovered that distant residue pairs, which move further away from each other in K-Ras^G12D^ also move away from their proximal neighbors. On the other hand, residue pairs that move closer to each other in K-Ras^G12D^ assume closer conformations relative to their proximal neighbors.

### G12D mutation leads to increased fluctuations of loop residues in SII region

Next, to understand how the flexibility of K-Ras changes upon G12D mutation, we calculated the root-mean-square fluctuations (RMSF) of each residue in both wild-type and mutant protein, where RMSF is a measure of the average atomic fluctuations of a residue.

First, we investigated the effects of the mutation on active wild-type and mutant protein flexibilities, and observed that in the mutant protein, the fluctuations of the loop residues of SII are increased (Fig 6A). Our residue pair distance calculations in active mutant K-Ras also showed that the residues in SII loop move away from some of the residues in α3 helix, and distances between those residue pairs display broad distributions (Fig 3). Considering the increased fluctuations of SII, these broad distance distributions with larger peak values between SII and the other parts of the protein may be arising from the increased flexibility of SII due to G12D mutation.

**Figure 6.**
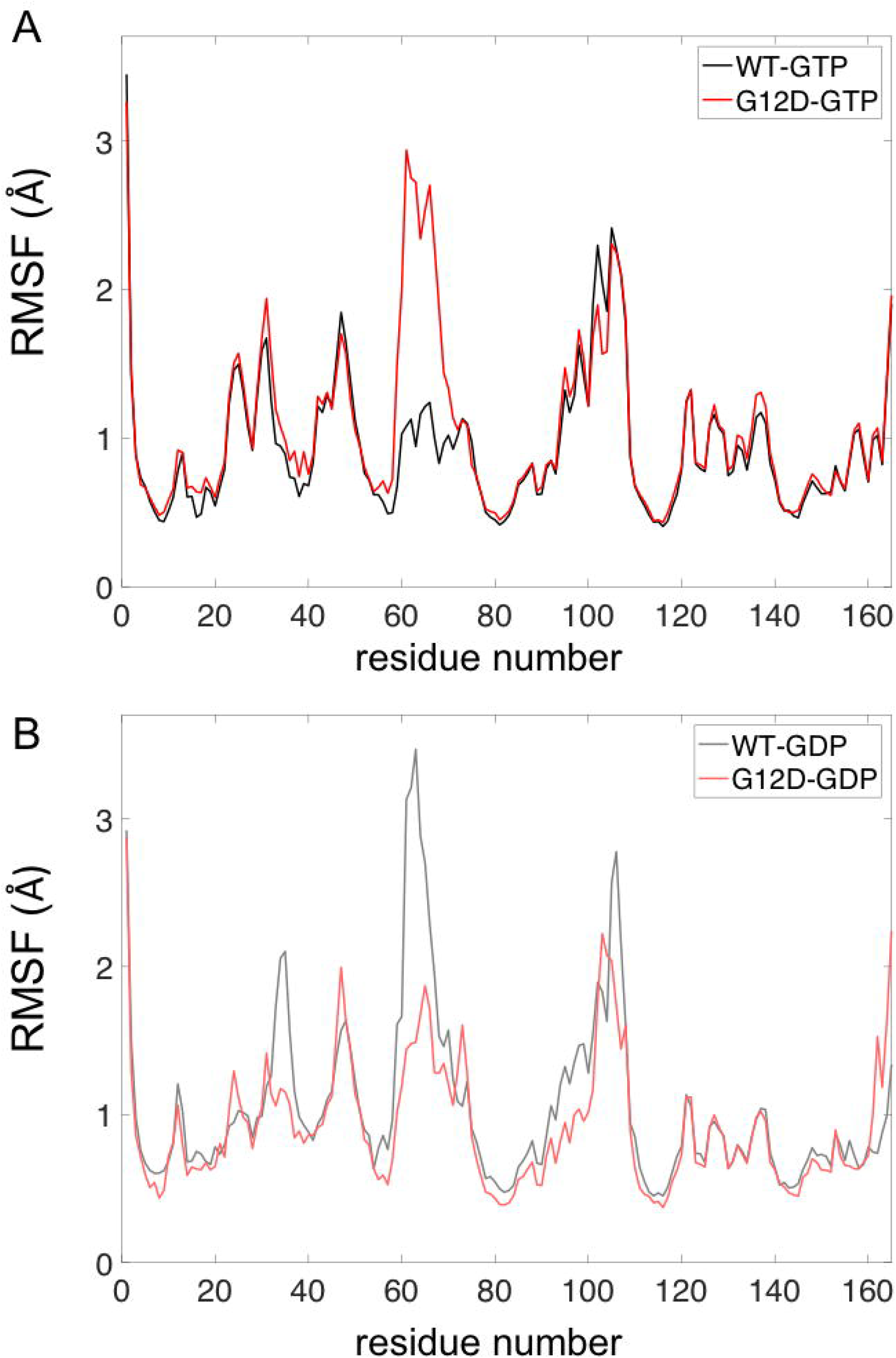
Dynamic changes in K-Ras upon G12D mutation. (A) RMSF values of K-Ras^WT^-GTP (black) and K-Ras^G12D^-GTP (red) residues. (B) RMSF values of K-Ras^WT^-GDP (grey) and K-Ras^G12D^-GDP (pink) residues.

We then calculated the effects of mutation on inactive K-Ras flexibility by comparing residue fluctuations of wild type and mutant protein. Figure 6B shows that the effect of G12D mutation on inactive (GDP-bound) protein is opposite to its effect on the active (GTP-bound) form, where the SII loop residue fluctuations decrease after G12D mutation in K-Ras–GDP. Furthermore, SI fluctuations also decreased in mutant K-Ras–GDP.

### G12D mutation leads to increased SII loop fluctuations in the active protein, making it unable to form salt bridges

From the analysis of the MD simulation datasets, we observed that E63 (SII) forms a salt bridge with R68 (SII) in active form in wild-type K-Ras, but not in active mutant. The increased fluctuations of the SII loop in the absence of E63-R68 bond are shown in Figure 6A. However, in inactive mutant, this salt bridge is intact and the fluctuations of the SII loop are decreased, as shown in Figure 6B.

### G12D mutation markedly increases the negatively correlated motions of SII residues in active K-Ras

Regulation of protein dynamics is strictly coordinated by the correlations of residue fluctuations. Figure 7 presents pairwise correlations of residue fluctuations (*C_ij_*), for K-Ras^WT^-GTP (upper-left), K-Ras^G12D^-GTP (upper-right), K-Ras^WT^-GDP (lower-left) and K-Ras^G12D^-GDP (lower-right). In K-Ras^WT^-GTP, β2-β3 regions move in negative correlation with α1-SI region and β6; but this correlation is disturbed by the G12D mutation. However, the most striking effect of the mutation on the active protein is the occurrence of new correlations between the motions of the SII loop and other protein regions. As seen in Figure 7B, upon G12D mutation, motions of residues in SII become negatively correlated with those of residues in P-loop, β3, β4 and α3-Loop7. For the inactive GDP-bound form, G12D mutation does not have a notable effect on motion correlations.

**Figure 7.**
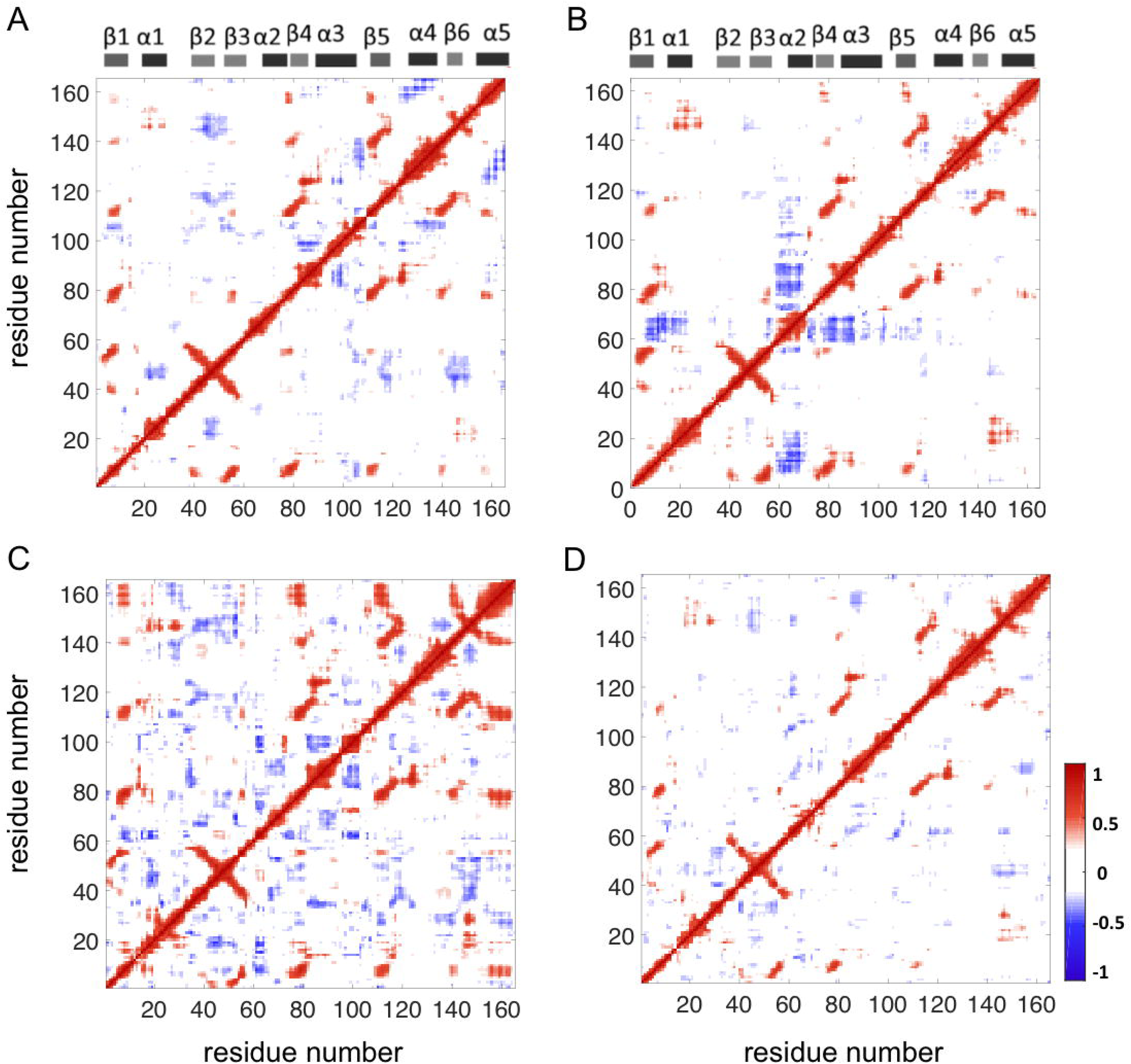
The correlated residue motions in K-Ras. Correlation coefficient maps for K-Ras^WT^-GTP (A), K-Ras^G12D^-GTP (B), K-Ras^WT^-GDP (C) and K-Ras^G12D^-GDP (D). Red dots show positive correlations and blue dots show negative correlations.

### G12D mutation does not affect the coupled motions of the central β-sheet

As seen in Figure 1, K-Ras structure consists of a central β-sheet that is composed of six β-strands, which are surrounded by five α-helices. Our correlation calculations show that the β-strands fluctuate in correlation in both the active and inactive wild type forms (Fig 7A, C). Specifically, motions of β1 are coupled to motions of the other strands; β2 and β3 motions are coupled to each other and motions of β5 are coupled to motions of β4 and β6. These coupled motions within the central β-sheet are not altered by the G12D mutation.

### G12D mutation causes negatively correlated fluctuations in regions (Fig 7B) that move away from each other in inactive protein

We observed that after G12D mutation, negative correlations occur between the regions -SII-P-loop and SII-α3- which move away from each other (Fig 2A) and from their neighbors (Fig 2B). The combination of distance (Fig 2) and correlation (Fig 7B) calculations reveals the relation between conformational and dynamic changes in the protein as a result of the G12D mutation.

## Discussion

K-Ras is an important GTPase in cellular signaling that is only active in its GTP-bound state ^8,9^. Structurally, in wild-type active K-Ras, the P-loop, SI and SII switch regions are bound to the phosphate groups of GTP and are responsible for its GTPase function. However, when there is a G12D mutation in the P-loop, GTP hydrolysis is impaired and K-Ras freezes in its active state ^9^, causing uncontrollable cellular growth and evasion of apoptotic signals ^27,48,49^. Despite extensive literature on the effects of G12D mutation on K-Ras, its effects on the regulation of local dynamics of the active and inactive protein, and how these relate to its effects on structure and local conformations of the protein remain unknown. At the same time, since protein function is intrinsically related to its dynamics, this information can support studies on effective targeting of mutant K-Ras. To understand the local changes in the dynamic behavior of K-Ras caused by the G12D mutation, we first identified the structural and conformational changes in its residues and then related them to changes its dynamic characteristics by MD simulation data analysis of both wild type and mutant K-Ras in GTP- and GDP-bound forms.

Using residue pair distance analysis, we have discovered that K-Ras accesses a range of conformational states during its simulations, and that the G12D mutation alters the distribution of its conformational states, which is specific to the bound nucleotide. Overall, the residue pair conformational states, which we present in Figures 3-5, populate multiple distances in inactive (GDP-bound) wild-type and mutant K-Ras. However, they adopt two different conformational states in active (GTP-bound) wild-type protein and only a single predominant conformational state in active mutant protein. Specifically, the residue pairs between SII-α3 have two different conformational states, which we have labeled as first and second conformational states in *Results*. While the wild type protein populates both conformational states, the second conformation is predominant in the G12D mutant. In the first conformation, the N-terminus of SII is positioned toward N-terminus of α3 helix (Fig 3), and the central residues of SII move away from C terminal end of α3 (Fig 5C). In this conformation, we observed an H-bond network within SII residues: (i) R68 forms bonds with E63, S65, Y71, M72, (ii) M67 is bound to Q70 and (iii) Q61 is bound to E63. However, in the second conformation, the central residues of SII move toward C terminal end of α3, this H-bond network disappears and SII forms H-bonds with α3.

Our observation of two conformational states for active wild-type K-Ras is consistent with previous experimental studies ^50-52^, which have shown that the protein has two states, a catalytically incompetent (T) and a catalytically active (R) state, and that SII can assume a variety of positions due to the conformations of α3 helix and loop L7 that associate with these states (please note that as shown in Figure 1, L7 is right adjacent to α3, where α3 ends at residue 103 and the loop begins at residue 104). Briefly, these studies have shown that in “T” state, α3-L7 moves toward SII and disrupts the H-bond network centered at R68 in SII ^53^. Downstream signaling remains ‘on’, as GTP hydrolysis with disordered SII region is slow. On the other hand, in “R” state, H-bond network in SII is intact, allowing for catalytic activity and turning K-Ras signaling “off”. Our results provide an atomistic level structural explanation to these observations by revealing that G12D mutation shifts the two SII-α3 conformations in favor of a single predominant conformation (the second conformation), disrupting the H-bond network between SII residues (Fig 3). In summary, our results suggest that the deviation of central SII residues toward α3 helix, and the disruption of the H-bond network within SII may impair GTP hydrolysis, leading to the constitutive activation of K-Ras^G12D^–GTP signaling.

In our residue pair distance analyses, we also observed that in wild-type K-Ras-GTP, the distribution curves of the distances between the P-loop and Q61 are Gaussian-shaped, narrow dispersion curves. However, in G12D mutant K-Ras-GTP, they significantly deviate from the Gaussian. Since Q61 is a known critical catalytic residue for both intrinsic and GAP-mediated GTP hydrolysis, it is possible that this highly variable nature of the P-loop-Q61 distance, suggesting high flexibility, affects GTP hydrolysis in K-Ras^G12D 10^. Furthermore, using the average of residue pair distances for each residue, 〈Δ*R̄i*〉, we observed that the P-loop, SI, SII and α3 regions move away from their neighbors upon G12D mutation (Fig 2B).

In the MD simulations, we observed that K-Ras G12D mutation further alters the H-bond network and formation of salt bridges. Because of the formation and breakage of the bonds between different parts of the protein, the P-loop, SII and α3 regions of mutant K-Ras assume conformations different than the wild-type. Furthermore, by correlating the conformational and dynamic changes in residues at these regions, we discovered that the G12D mutation leads to the coupling of the motions of SII region with those of the residues at these sites. The effects of the G12D mutation in increasing these correlations are clearly observed in pairwise correlation maps (Fig 7B), which show marked differences between K-Ras^G12D^ and K-Ras^WT^, and are also consistent with the increased amplitude of SII fluctuations, as shown in Figure 6. Our results are consistent with a previous computational study that showed that SII displays increased fluctuations and negative correlations with other parts of the protein ^27^. Consequently, we show that the G12D mutation leads to characteristic population of relative SII conformations and also changes in SII flexibility and dynamics. Our study goes beyond this by identifying the atomistic basis of the changes in dynamic behavior of K-Ras in terms of bond formations.

Overall, these results provide a new understanding of the local changes in dynamic behavior of K-Ras^G12D^ and an atomistic basis for this behavior. Such an understanding can support studies that target the protein with small molecules, which can be an effective strategy for the ever elusive allosteric inhibition of oncogenic K-Ras^G12D^.

## Methods

### MD Simulations

We performed all-atom MD simulations for both GTP- and GDP-bound forms of K-Ras^WT^ and K-Ras^G12D^. We obtained the K-Ras-GTP^WT^ and K-Ras-GDP^WT^ structures from the final frame of the simulations of active and inactive states proteins by Vatansever *et al*, respectively ^54^. For constructing K-Ras^G12D^ structure, we mutated glycine to aspartate at position 12 in K-Ras-GTP^WT^ structure using Discovery Studio 4.5 software, (DS) ^55^. To optimize the K-Ras^G12D^-nucleotide complex, we used Clean Geometry tool of DS. For MD simulations, we used NAMD 2.10 ^56^ with AMBER ff99SB ^57^ and general amber force fields (GAFF) ^58^. Briefly, we performed energy minimization of the initial model after we introduced the G12D mutation in K-Ras, and then ran MD simulations of each complex following the protocols from Vatansever *et al* ^54^, the details of which we provide in Supplementary Methods. During the simulations, we applied minimization for 10,000 steps and equilibration for 500,000 steps, after which we performed 1 microsecond MD simulations, and saved atomic coordinates *R̂* of all atoms every 10ps. We repeat the same simulation steps for two independent simulations of each complex. We used the last 900ns of the simulation trajectories in all computations described in this study. To eliminate all rotational and translational motions, we aligned the trajectories to the first frame using VMD software 1.9.2 ^59^. We visualized the trajectories with VMD. To identify salt bridges formed in the protein during the MD simulations, we used Salt Bridges Plugin, Version 1.1, of VMD.

### Principal component analysis and projection

Consider a protein with N residues (for K-Ras protein N=165), where the position vector for the *i* ^th^ atom, *r_i_* = [*x_i_ y_i_ z_i_*], is known from simulations performed for *M* time steps. First, we mean-center the coordinate vector by subtracting the temporal averages from each of its coordinates, i.e.,

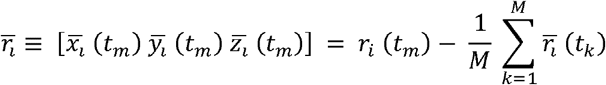

Then, we calculate the magnitude (|| ||) of the coordinate vector:

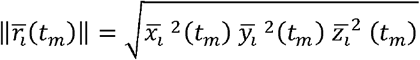

and represent the MD simulation data performed for *M* time steps in an *N*×*M* matrix:

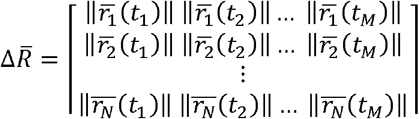

Then, we calculate the covariance matrix (*C*), which is an *N*×*N* symmetry matrix, by taking the dot product of ∆*R̄* and ∆. *R̄*^T^ as below:

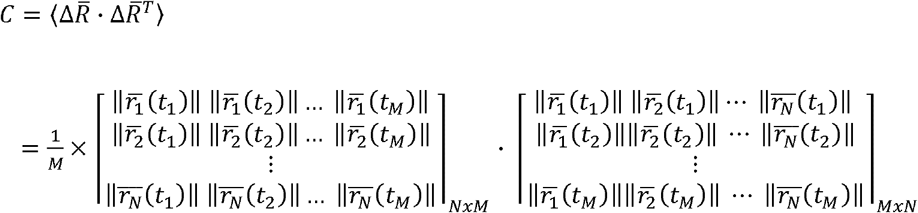

Therefore, each element of the covariance matrix, *C_ij_* is computed from:

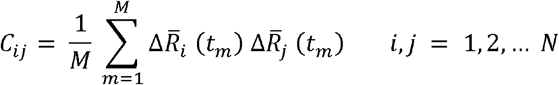

where ∆*R̄_i_*(*t_m_*) denotes the *i^th^* component of ∆*R̄*(*t_m_*). Since, is a symmetry matrix, it can be diagonalized with a symmetric eigenvalue decomposition as, VΛV^T^, where V= (V_1_, V_2_, …, V_N_) is a set of real-valued eigenvectors, Λ is a real diagonal matrix, and the eigenvector V_*i*_ corresponds to the eigenvalue Λ_*ii*_ and the *i^th^* column vector of V is the *i^th^* PCA mode. The projection of simulation data ∆*R̄*(*t_m_*) m onto the *i*^th^ PCA mode is computed from:

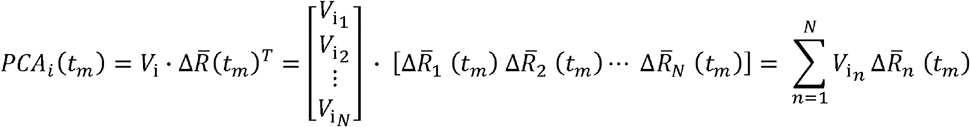

Iterating this *PCA_i_*(*t_m_*) computation over *m* for all *M* time intervals, we obtain the time series *PCA_i_* = [*PCA_i_*(*t*_1_) *PCA_i_*(*t*_2_) … *PCA_i_*(*t_M_*)]for *i^th^* mode. Specifically, we calculated the first (PCA_1_) and the second (PCA_2_) modes for projection of MD trajectories onto principle components (Supplementary Figure S2.). Based on the PCA projection plots, we separated the *M* time points of the trajectories into PCA states, determined the most densely populated states and calculated the probabilities of these states. In calculations of the residue pair distances and correlations, we used the trajectory points in the most densely populated PCA states and weighted the calculations by the probability of its state.

### Residue pair distance calculations

To quantify the effect of the G12D mutation on the distances between K-Ras residue pairs, we developed a new computational algorithm detailed in Supplementary Figure S1. Briefly, we first assumed K-Ras^WT^ as the initial state and K-Ras^G12D^ as the final state. Then, we calculated the distances between Cα atoms of two residues (*i*, *j)* as we previously described ^54^ and listed in Supplementary Methods. Then we borrowed the ‘first coordination shell’ definition from the Gaussian network model (GNM), which is widely used in the analysis of protein dynamics. Studies that use GNM typically assume the ‘first coordination shell’ as the maximum Cα-Cα distance for the separation between two contacting residues at ∼7.2Å ^44,45 46,47^. We followed this protocol and determined the first coordination shell around a selected residue by choosing its Cα as the center of a volume V with a radius of r1 ∼7.2 Å^45^. However, because the contribution of non-bonded pairs to higher-order coordination shells may also be significant ^47,60^, we also studied residue pairs that are within their ‘second coordination shell’ in K-Ras^WT^ structure, which we defined at twice the volume of the first, with a radius of ∼9.1 Å ^47^.

For every residue pair *(i, j)* where *j* is in the second coordination shell of *i*, we first calculated its time-averaged distance in K-Ras^WT^ (*R̄_ij_* _WT_) and in K-Ras^G12D^ (*R̄_ij_* _G12D_). We then calculated the difference (∆*R̄_ij_*) between *R̄_ij_* _WT_ and *R̄_ij_* _G12D_, where ∆ *R̄ij* = *R̄_ij_* _G12D_ - *R̄_ij_* _WT_. The magnitude of the difference is the degree of distortion resulting from the G12D mutation. We present ∆ *R̄ij*, values in the pairwise distances map (Fig 2), where a positive value indicates that a residue pair moves apart upon G12D mutation, while a negative value indicates that the pair gets closer. Then, to identify all residue pairs (*ij*) that were significantly distorted by the G12D mutation, we selected the residue pairs that have the greatest (positive and negative) ∆ *R̄ij*, values. We assumed that the residue pairs with ∆ *R̄ij* >2.50 or < −1.40 showed the most significant distance changes. For those identified residue pairs, we plot the distribution graphs *W*(*R_ij_*) of their distances (*R_ij_*) during the simulations of K-Ras^WT^ and K-Ras^G12D^ (For details see Supplementary Methods).

Then, to quantify the changes in local volumes upon G12D mutation, for each residue *i* we calculated the average of all ∆*R̄_ij_* values based on the formula 〈∆*R̄i*〉 = Σ_*j*_∆*R̄ij*/*N_n_* where *N_n_* is the number of residues *j* in the second coordination shell of residue *i*. In detail, for a residue *i*, at the center of a volume V with a radius of 9.1 Å (the second coordination shell) and we defined the residues *j* within this volume V as the *neighbors* of residue *i*. Then, we calculated the total change in the distance between residue *i* and its neighbors, Σ_*j*_∆*R̄ij*, and divided it by the number of neighbors Σ_*j*_∆*R̄ij*/*N_n_*. The resulting 〈∆*R̄i*〉 value is a measure of the change in volume around residue *i* due to G12D mutation.

### Residue pair correlation calculations

To investigate the coupled motions of residue pairs in protein dynamics, we calculated the correlation coefficients between their fluctuations (*C_ij_*). A correlation coefficient value of a residue pair ranges from −1 to 1, where for residue pair fluctuations that are not coupled *C_ij_*= 0; perfectly positively correlated C_*ij*_=1, and perfectly negatively correlated C_*ij*_=−1. We calculated the correlation coefficients as described in our previous study ^54^, which is summarized in detail in Supplementary Methods.

The datasets generated during and/or analysed during the current study are available from the corresponding author on reasonable request.

## Acknowledgements

ZHG acknowledges funding from from the LUNGevity Foundation and start-up funds from the Icahn Institute at Icahn School of Medicine at Mount Sinai. We would also like to thank Dr. Roman Osman for valuable comments on the manuscript and Dr. Myvizhi Esai Selvan for help with one of the figures.

## Author Contributions

SV conducted the molecular dynamics simulations and data analyses, prepared the figures and evaluated results. ZHG and BE contributed to study design and evaluation of the results. ZHG and BE contributed oversight to the study. All authors have contributed to manuscript drafting and agree to the submitted manuscript.

## Competing Interests

The authors declare no competing interests.

